# Droplet-based microfluidic analysis and screening of single plant cells

**DOI:** 10.1101/199992

**Authors:** Ziyi Yu, Christian R. Boehm, Julian M. Hibberd, Chris Abell, Jim Haseloff, Steven J. Burgess, Ivan Reyna-Llorens

## Abstract

Droplet-based microfluidics has been used to facilitate high throughput analysis of individual prokaryote and mammalian cells. However, there is a scarcity of similar workflows applicable to rapid phenotyping of plant systems. We report on-chip encapsulation and analysis of protoplasts isolated from the emergent plant model *Marchantia polymorpha* at processing rates of >100,000 protoplasts per hour. We use our microfluidic system to quantify the stochastic properties of a heat-inducible promoter across a population of transgenic protoplasts to demonstrate that it has the potential to assess gene expression activity in response to environmental conditions. We further demonstrate on-chip sorting of droplets containing YFP-expressing protoplasts from wild type cells using dielectrophoresis force. This work opens the door to droplet-based microfluidic analysis of plant cells for applications ranging from high-throughput characterisation of DNA parts to single-cell genomics.

## Introduction

In light of recent advances in DNA synthesis and construct assembly, phenotyping of genetic circuits generated by these components is likely to soon limit the rate of scientific progress. This is particularly true for plant science, where the time required for generation of transgenic organisms ranges from months to years. Protoplasts, individual cells whose wall has been removed through mechanical or enzymatic means, offer an alternative to analysis of plant tissues and open up the possibility of high-throughput phenotyping of single cells^1^. Introduction of DNA into protoplasts by electroporation^2–7^, PEG-based transfection^8,9^, or particle bombardment^10^ has proven a valuable approach to transient and stable transformation of nuclear and organellular genomes, in particular for plants not amenable to *Agrobacterium*-mediated transgene delivery. Protoplasts have furthermore been used to overcome barriers of sexual incompatibility in generating hybrid plants with novel properties^11^. Following transformation or somatic hybridization, whole plants can be regenerated from individual protoplasts through tissue culture^12^. In addition, protoplasts have become recognized as convenient experimental systems for studying aspects of plant cell ultrastructure, genetics, and physiology^13^. However, to date protoplasts have been extracted and analysed in bulk, limiting their use. Recently, droplet-based microfluidics has gained increasing popularity as a platform for high-throughput culture, manipulation, sorting, and analysis of up to millions of individual cells under diverse conditions^14–18^.

This approach is based on pico-to nanoliter-volume aqueous microdroplets which spatially separate individual cells from one another during processing. To date, droplet-based microfluidics has primarily been applied to bacteria^19–23^, unicellular eukaryotes^23–25^, and nonadhesive mammalian cells^26–28^. The prospect of utilizing this platform for characterization and screening of individual plant protoplasts is highly attractive: high-throughput screening of whole plants is substantially limited by their slow growth and size. By contrast, millions of plant protoplasts may be processed in a matter of hours using droplet-based microfluidics, and so could allow pre-selected protoplasts to be regenerated into whole plants.

Microfluidic devices have been applied for the collection and lysis^29^, culture^30^, chemically-induced fusion^31^, electrofusion,^32^ regeneration^33^, and developmental characterization^34^ of plant protoplasts. However, no system for the high-throughput characterization or sorting of individual plants protoplast based on their level of gene expression has been reported to date. While widely used for cell sorting, FACS cannot currently be applied to plant protoplasts as their fragility causes them to rupture under strong acceleration. One group has thus used optical tweezers to displace non-encapsulated plant protoplasts in a microfluidic chip, but has not demonstrated successful sorting^35^.

In this paper, based on the genetic expression of a fluorescent reporter protein we demonstrate high-throughput characterization and sorting of plant protoplasts encapsulated individually in aqueous microdroplets. We use protoplasts derived from the model plant *Marchantia polymorpha*^36^, which combines a simple genomic structure^37^ with ease of handling^38^ and robustness of regeneration in absence of supplemented plant hormones^39^. We enzymatically isolate *Marchantia* protoplasts from adult thalli, and encapsulate them via a flow-focusing microfluidic device. An optical detection setup integrated into the microfluidic channel allows high-throughput quantification of chlorophyll autofluorescence or promoter-controlled YFP fluorescence emitted by individual encapsulated protoplasts. We demonstrate how this droplet-based microfluidic system can be used to rapidly measure the stochastic properties of an inducible plant promoter over a population of individual plant protoplasts. We furthermore show this system is capable of automated sorting of individual encapsulated protoplasts based on their YFP fluorescence intensity. Facilitating high-throughput screening and enrichment of plant protoplasts based on expression of a fluorescent reporter gene, our microfluidic system streamlines the identification and isolation of desired genetic events in plant biology research and modern biotechnology.

## Results and discussion

Isolated *Marchantia* protoplasts were encapsulated in microdroplets on a flow-focusing microfluidic device (Fig. 1A). The aqueous protoplasts suspension flowed perpendicularly to two streams of fluorinated carrier oil containing PicoSurf1 non-ionic surfactant. The two phases intersected at the ‘flow-focusing junction’, as the oil streams enveloped the droplet that budded off from the aqueous stream (Fig. 1B). The density of *Marchantia* protoplasts was adjusted to ensure microdroplets contained no more than one protoplast each (Fig. 1C), which is important for accurate quantification of cellular fluorescence intensity. The same approach was also successful for encapsulation of the widely used angiosperm model *Arabidopsis thaliana* (S1, S2 ESI†).

**Figure 1.**
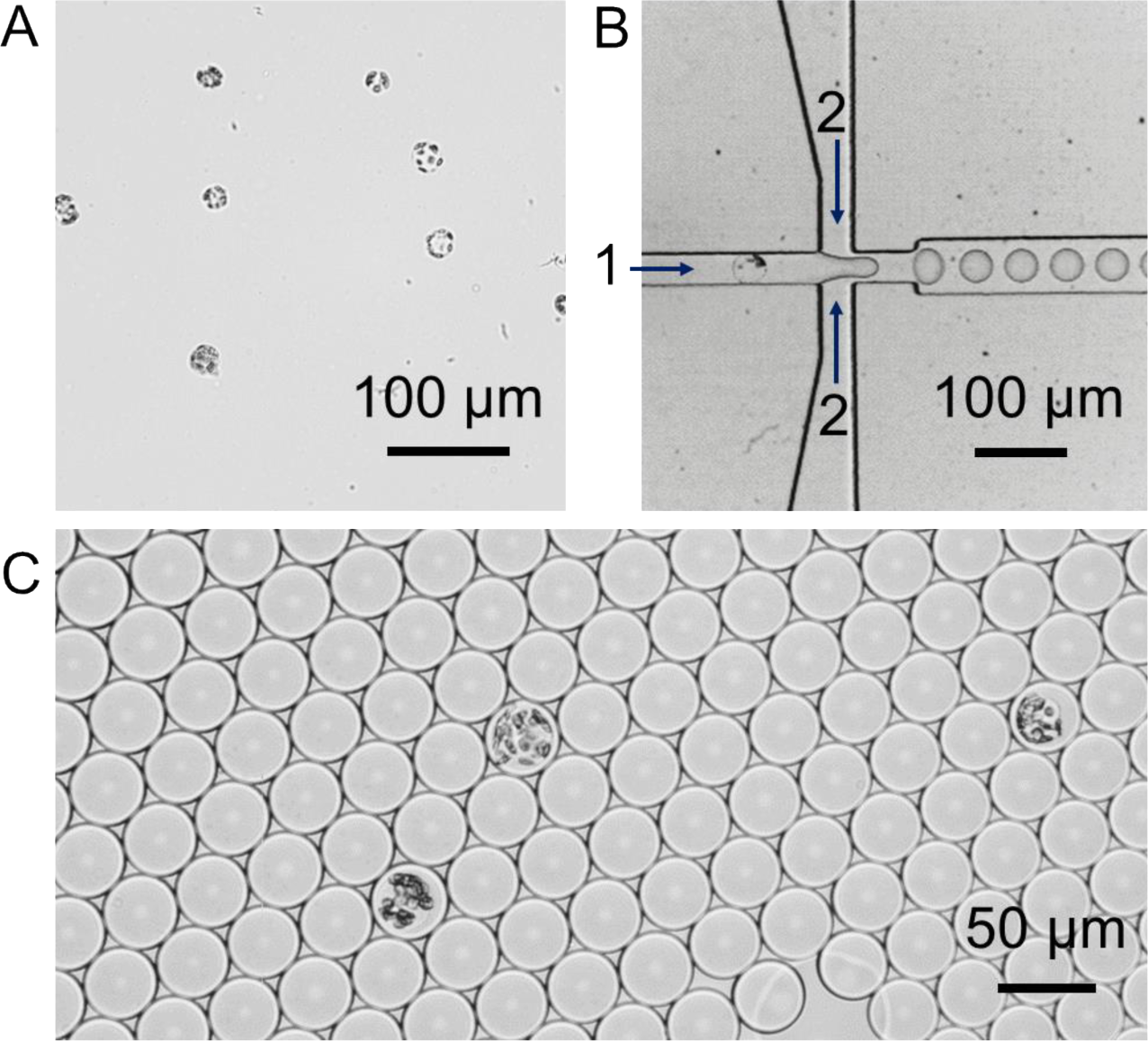
(A) Bright field micrograph of *Marchantia* protoplasts isolated from mature thalli. (B) Bright field micrograph of a flow-focusing microfluidic device for encapsulation of *Marchantia* protoplasts in water-in-oil microdroplets. (C) Bright field micrograph of individual *Marchantia* protoplast encapsulated in microdroplets.

While encapsulated, protoplasts remained intact over a period of at least 12 hours (Fig. 2). To quantify chlorophyll autofluorescence in individual encapsulated protoplasts, an optical setup was integrated to the system (Fig. 3). Each microdroplet was re-injected into a microfluidic flow channel continuously exposed to a 491 nm laser beam. Fluorescence emitted from excited protoplasts passed through a 633 nm longpass filter and the signal was collected by a photomultiplier tube (PMT). Using this experimental approach, the fluorescence of each protoplast was quantified reaching a potential rate of 115,200 individual protoplasts per hour. This observation suggests that high-throughput quantification of chlorophyll fluorescence using our microfluidic setup can be utilized for assessment of the quality of a protoplast preparation. The same experimental approach was also used for quantification of reporter protein fluorescence in individual plant cells, as illustrated by protoplasts derived from transgenic mpt0 *M. polymorpha* constitutively expressing mVenus_40_ yellow fluorescent protein (YFP) under control of the strong constitutive MpEF1α promoter_41_ (Fig. 4).

**Figure 2.**
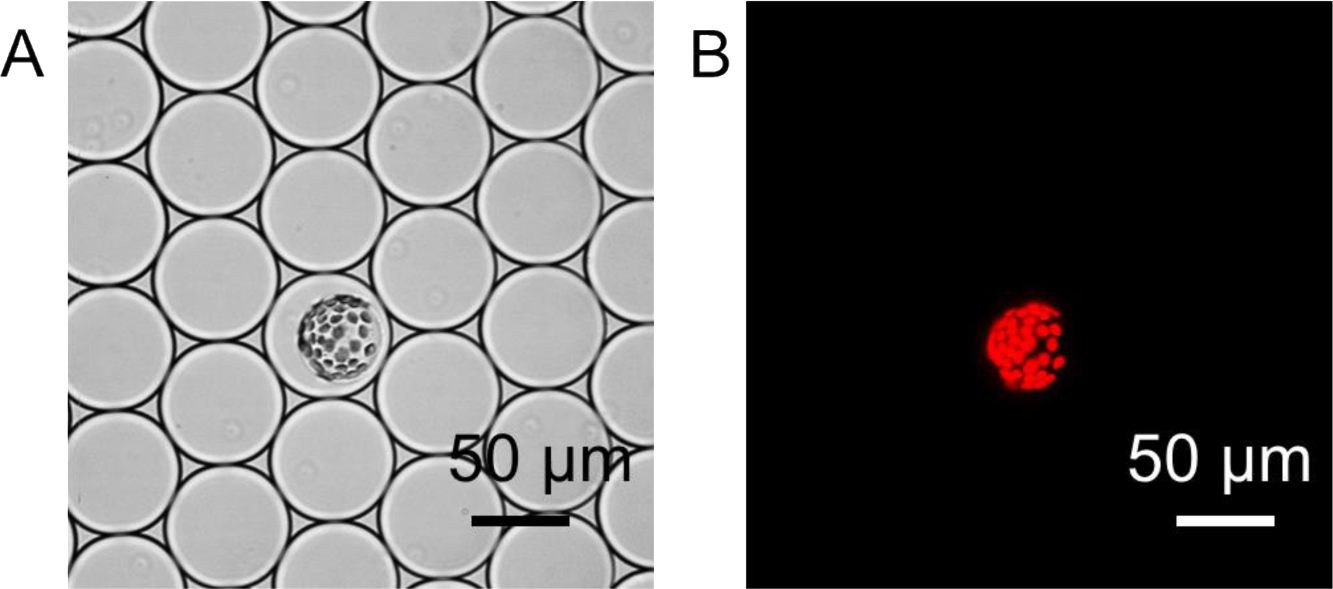
(A) Bright field and (B) chlorophyll fluorescence micrographs of individual *M. polymorpha* protoplast after 12 hours of encapsulation in microdroplets.

**Figure 3.**
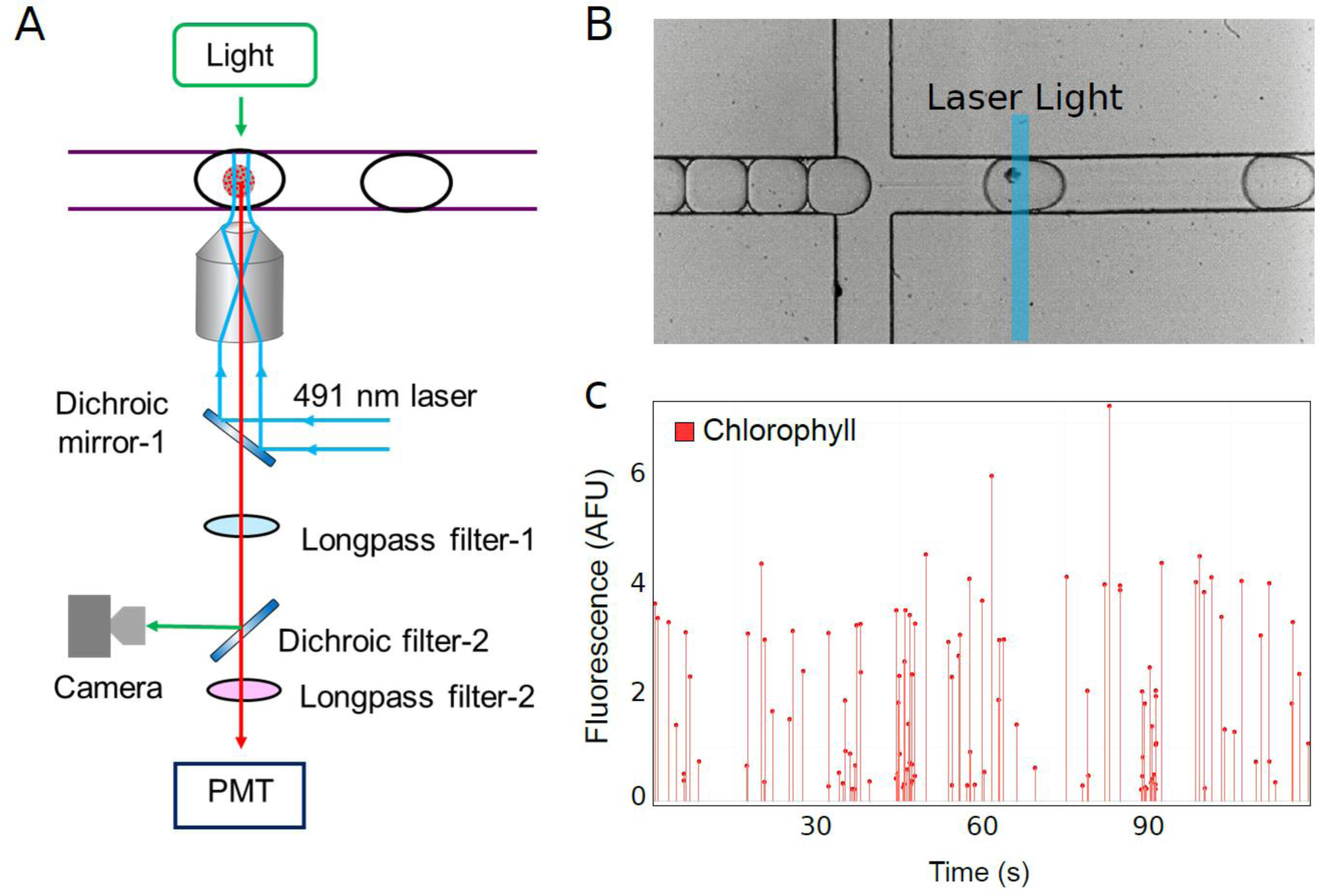
(A) Experimental setup used for quantification of fluorescence intensity of encapsulated protoplasts. Long-pass filter-1: 495 nm; long-pass filter-2: 633 nm; dichroic filter-1: 495 nm; dichroic filter-2: 633 nm. (B) Bright field micrograph of an encapsulated protoplast passing through the excitation laser beam. (C) Representative photomultiplier tube (PMT) readout of chlorophyll fluorescence intensity represented as arbitrary fluorescent units (AFU) recorded on chip over 120 s. Each line represents an individual encapsulated protoplast.

**Figure 4.**
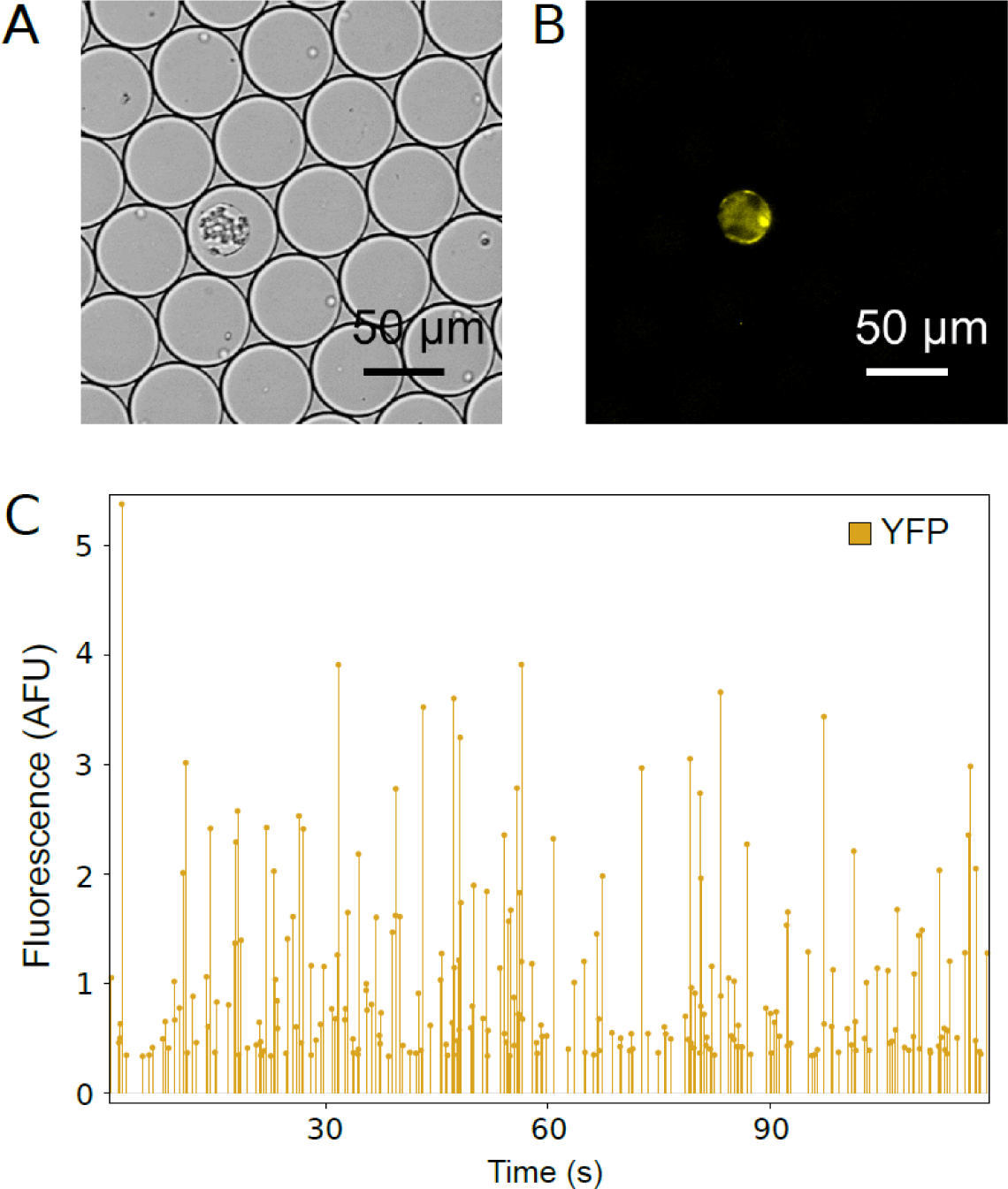
(A) Bright field and (B) YFP fluorescence micrograph of an individual encapsulated protoplast derived from transgenic mpt0 *M. polymorpha* constitutively expressing mVenus YFP. (C) Representative photomultiplier tube (PMT) readout of YFP fluorescence intensity represented as arbitrary fluorescent units (AFU) recorded on chip over 120 s. Each line represents an individual encapsulated protoplast.

As the next step, our system was applied for the analysis of the stochastic activity of an inducible promoter across a population of individual plant cells. For this purpose, transgenic PMpHSP17.8 lines of *M. polymorpha* were generated, which expressed mVenus yellow fluorescent protein (YFP)_40_ under control the endogenous heat-responsive MpHSP17.8 promoter. It was previously shown that incubation of transgenic *M. polymorpha* at 37°C for 1 h induced a P_MpHSP17.8_-controlled targeted gene by approximately 700-fold^42^.

To measure the stochastic properties of this promoter, transgenic PMpHSP17.8 *M. polymorpha* was incubated under two different temperature conditions and isolated protoplasts from each sample for on-chip quantification of YFP fluorescence (Fig. 5). *M. polymorpha* thalli were either subjected to (i) 2 h at 37°C followed by 2 h at room temperature or to (ii) 4 h at room temperature (control). Protoplasts isolated from heat-shocked plants exhibited significantly higher levels of YFP activity compared to the Control (p < 2.2e-16, 95% CI [-0.2, -0.13]. This result illustrates the power of our microfluidic system to quantify stochastic properties of plant promoters as a function of environmental conditions. An even more powerful application of our microfluidic platform is sorting of individual encapsulated protoplasts based on their level of expression of a target reporter gene. This allows single plant cells to be pre-screened for downstream sequencing and/or regeneration of whole plants. For this purpose, a microdroplet-based microfluidic sorting system was developed (Fig. 6A): two oil flow-focusing channels allowed the spacing between microdroplet to be controlled by flow-rate adjustment. Microdroplet sorting was implemented by a pair of electrodes generating a dielectrophoretic force applied to the microdroplet. When the electrodes were off, the microdroplets were pushed into the “negative” channel due to its lower fluidic resistance compared to the “positive” channel. Switching the electrodes on steered the individual microdroplets into the “positive” channel through dielectrophoretic force. The generation of an electrode pulse was dependent on the fluorescence intensity emitted from each microdoplet: microdroplets were steered to the “positive” channel only if they contained a protoplast expressing YFP above-threshold levels of 1.3 arbitrary fluorescence units (AFU; see video S3, ESI†). The platform was tested using microdroplets containing protoplasts isolated from either wild type or transgenic mpt0 *M. polymorpha*. Protoplast from both populations were pooled together and reinjected into the sorting device (Fig. 6B). Sorting successfully separated mVenus-expressing mpt0 protoplasts from wild type protoplasts (Fig. 6C). This result showed our microfluidic platform capable of high-throughput selection of desired events across large populations of genetically diverse individual plant cells.

**Figure 5.**
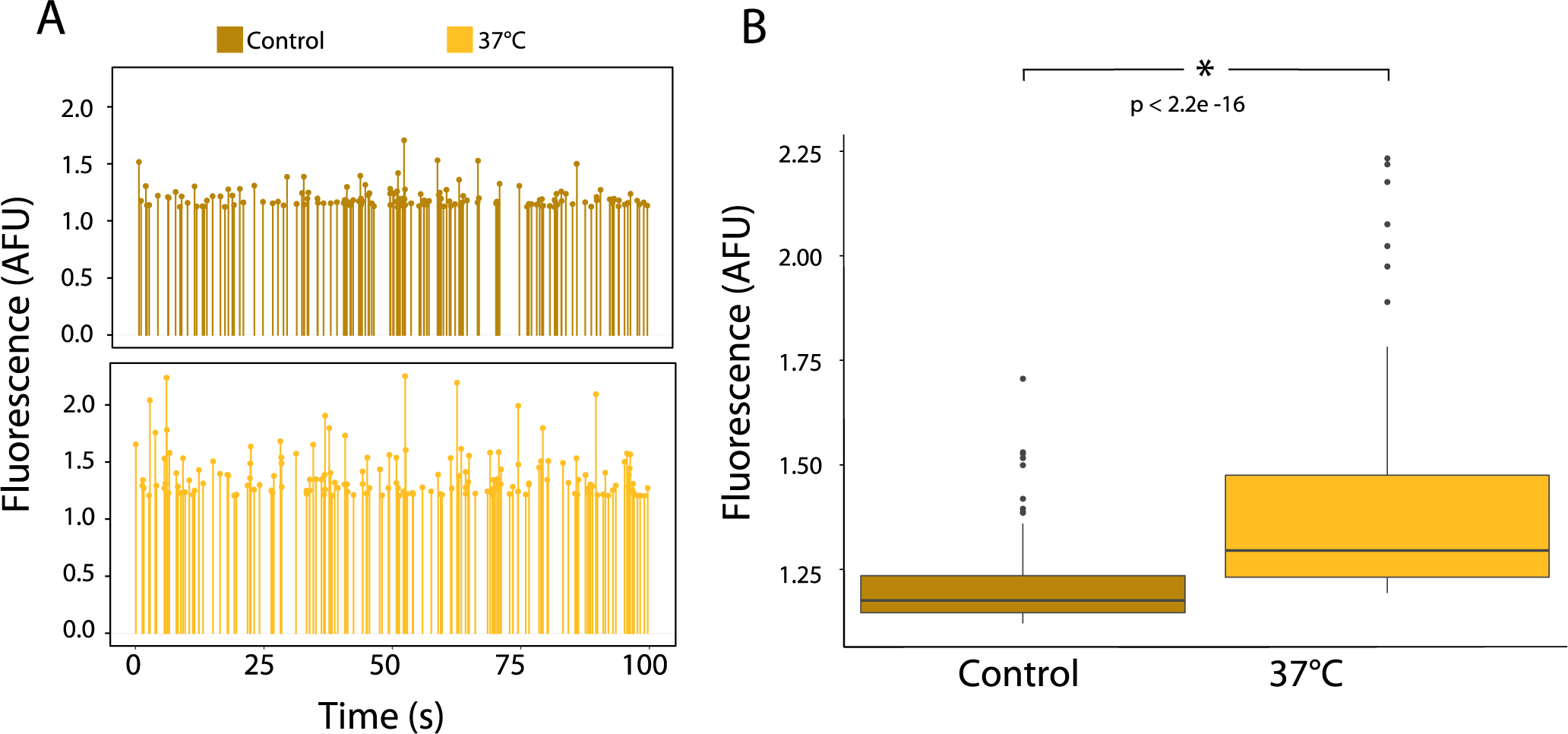
Characterization of heat-responsive induction of mVenus YFP in transgenic PMpHSP17.8 *M. polymorpha* in individual encapsulated protoplasts. Transgenic PMpHSP17.8 *M. polymorpha* encoding *mVenus* under control of the MpHSP17.8 promoter were either subjected to 4 h at room temperature (Control) or to 2 h at 37°C followed by 2 h at room temperature (37°C). Representative photomultiplier tube (PMT) readout of YFP fluorescence intensity represented as arbitrary fluorescent units (AFU) for protoplasts isolated from thalli subjected to either temperature treatment. Each line represents an individual encapsulated protoplast. (B) Boxplot of the difference in YFP fluorescence intensity between the two temperature treatments based on protoplast populations recorded on chip over 100 s each. The p value shown was calculated using unpaired t-test.

**Figure 6.**
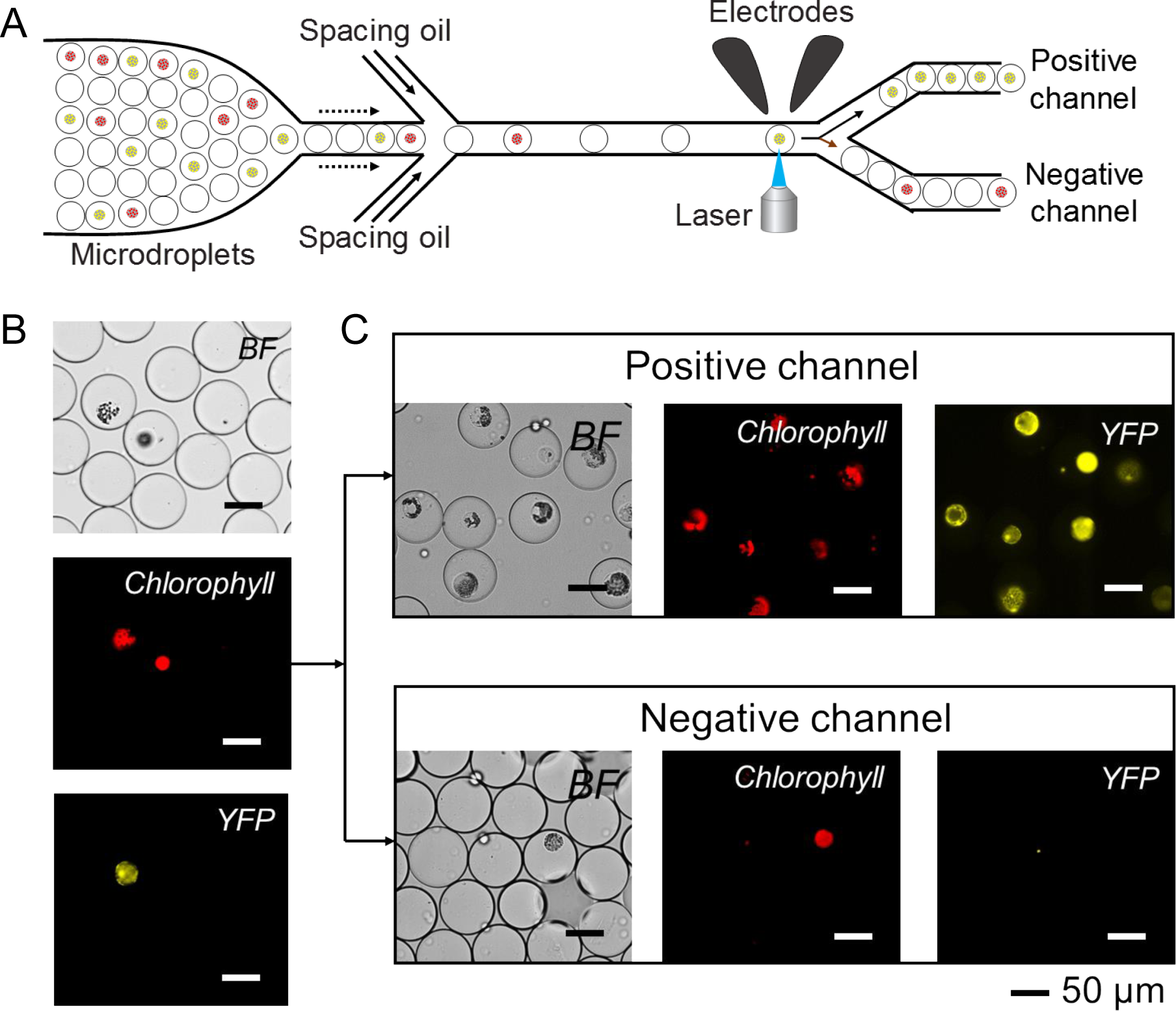
(A) Schematic representation of a platform for microfluidic sorting of encapsulated protoplasts. (B) Bright field and fluorescence micrographs of adjacent microdroplets containing protoplasts derived from wild-type and transgenic mpt0 *M. polymorpha*, respectively. (C) Bright field and fluorescence micrographs of microdroplets sorted into positive and negative channels based on their YFP fluorescence intensity.

## Materials and methods

### Chemicals, buffers, and media

Unless noted otherwise, chemicals used were obtained from Sigma Aldrich (Haverhill, UK) or Fischer Scientific (Loughborough, UK). DNA primers and Driselase from *Basidiomycetes sp.* (D8037) were obtained from Sigma Aldrich (Haverhill, UK). Standard molecular biology buffers and media were prepared as described in by Sambrook and Russell_43_.

### Microfluidic device fabrication

The microfluidic device was fabricated via soft lithography by pouring poly(dimethylsiloxane) (PDMS) along with crosslinker (Sylgard 184 elastomer kit, Dow Corning, Midland, MI, USA; pre-polymer: crosslinker = 10 : 1) onto a silicon wafer patterned with SU-8 photoresist_44,45_. The mixture was degassed in a vacuum dessicator and baked at 75°C overnight. The devices were peeled from the moulds and holes punched for inlets and outlets using a 1 mm diameter biopsy punch. The channel surface of PDMS was activated using oxygen plasma and attached to a glass slide. To ensure permanent bonding, the complete device was baked overnight at 110°C. The inner surface of the microchannels was rendered hydrophobic by flowing trifluorooctylethoxysilane through the channels, and the device was baked at 110°C for 2 h. Electrodes were incorporated into microfluidic chips by inserting a low-melting point indium alloy wire into a punched hole, and melting over a hot plate. Electrical wires were stripped at the end and inserted into the molten indium alloy (see also dx.doi.org/10.17504/protocols.io.ftybnpw).

### Binary vector construction

Binary vectors pCRB mpt0 (See Genbank accession No. MF939095) and pCRB PMpHSP17.8 (see Genbank accession No. MF929096) were based on pGreenII_46_, and constructed by means of isothermal assembly_47_. To confer hygromycin resistance to transgenic *M. polymorpha*, both binary vectors contained a hygromycin phosphotransferase gene_48_ expressed under control of the strong constitutive MpEF1α promoter_41_. pCRB further contained an *mVenus* yellow fluorescent reporter gene^40^ under control of P_MpEF1α_. pCRB PMpHSP17.8 contained an *mVenus* gene under control of the heat-inducible MpHSP17.8 promoter_42_.

### **Transformation of *A. tumefaciens***

50 μL aliquots of electrocompetent *A. tumefaciens* GV3101(pMP90) cells containing the pSoup helper plasmid were thawed on ice, mixed with 50-100 ng of DNA at the bottom of a pre-chilled 2 mm gap electroporation cuvette (VWR, Radnor, PA, USA), and kept on ice for 15 min. Electroporation was carried out using an *E. coli* Pulser Transformation Apparatus (Bio-Rad, Hercules, CA, USA) according to the manufacturer's instructions at 2.50 kV, 5 ms pulse length, and 400 ω default resistance. 1 mL of liquid SOC medium pre-warmed to 28°C was then immediately added to each cuvette, and the cells transferred to 15 mL Falcon tubes for recovery over 2-3 h at 28°C under shaking (ca. 120 rpm). 250 μL of cells were then spread onto LB 1.2%_(w/v)_ agar plates containing 25 μg/mL gentamicin, 5 μg/mL tetracycline, 50 μg/mL rifampicin, and 50 μg/mL kanamycin. Colonies became visible on the agar plates after approximately 2 days of incubation at 28°C.

### Plant materials and growth conditions

*M. polymorpha* Cam-strain plants were grown on B5 medium supplemented with 1.6 g/L vitamins (1/2 B5_Vit_; G0210, Melford, Ipswich, UK) containing 1.2%_(w/v)_ agar, under continuous white light.

### Surface sterilization and germination of *M. polymorpha* spores

*M. polymorpha* sporangia (2 per nuclear transformation to be attempted) were crushed with a polypropylene cell spreader until only small fragments (< 5 mm in diameter) remained visible. Sterile dH_2_O (1 mL per nuclear transformation) was added, and the tube vortexed vigorously for 30 sec. The crushing and vortexing steps were repeated, the suspension passed through a Falcon 40 μm cell strainer (Corning, Wiesbaden, Germany) to remove plant debris, and 500 μL aliquots of the filtrate transferred into 1.5 mL Eppendorf tubes. Spores were spun down at 13,000 rpm for 1 min, and the supernatant removed without disturbing the pellet. Each pellet was then resuspended into 1 mL of a sterilizing solution prepared by dissolving 1 Milton Mini Sterilizing Tablet (Procter & Gamble, Cincinnati, OH, USA) in 25 mL of sterile dH_2_O. The tubes were shaken at room temperature for 20 min at 200 rpm. Surface-sterilized spores were then pelleted by centrifugation as above and washed by 1 mL of sterile dH_2_O. The spore content of each tube was resuspended in 100 μL of sterile dH_2_O and spread on two 1/2 B5_Vit_ 1.2%_(w/v)_ agar plates. The plates were sealed and kept inverted under white fluorescent light at 23°C as described above. Small thalli were visible under a stereomicroscope after approximately 1 week.

### Nuclear transformation of *M. polymorpha* sporelings

2-3 colonies of *A. tumefaciens* GV3101(pMP90,pSoup) carrying a binary plasmid of interest were used to inoculate 4 mL of selective LB medium supplemented by 100 μM acetosyringone, and the culture incubated overnight at 28°C under shaking (ca. 120 rpm). 1 mL of the overnight culture was used to inoculate 4 mL of selective 1/2 B5_Vit_ medium supplemented by 100 μM acetosyringone, 0.1%_(w/v)_ casamino acids, 0.03%_(w/v)_ glutamine, and 2%_(w/v)_ sucrose (1/2 B5_VitAcSuc_). The diluted culture was incubated at 28°C for 4 h under shaking (ca. 120 rpm). Germinating spores of *M. polymorpha* on day 6 after surface sterilization were harvested by adding 2 mL of 1/2 B5_VitSuc_ (equals 1/2 B5_VitAcSuc_ without acetosyringone) to each plate, resuspending germinating spores in the liquid, through scraping them off the agar using a polypropylene cell spreader, and transferring the suspension to a 50 mL Falcon tube using a pipette. For each transformation, a suspension of germinating spores corresponding to the content of 2 agar plates (i.e. 2 sporangia) was diluted into 50 mL of 1/2 B5_VitAcSuc_ in a baffled 250 mL Erlenmeyer shaking flask. Following addition of 1 mL of transgenic *A. tumefaciens* GV3101(pMP90,pSoup), subcultured in 1/2 B5_VitAcSuc_ as described above, each flask was shaken at 150 rpm for 48 h under white fluorescent light at 23°C. After co-cultivation, spores were rescued by passing the suspension through a Falcon 40 μm cell strainer (Corning, Wiesbaden, Germany). Collected spores were washed by 200 mL of 100 μg/mL cefotaxime in sterile dH_2_O to remove *A. tumefaciens*, and spread on 1/2 B5_VitAcSuc_ 1.2%_(w/v)_ agar plates containing 100 μg/mL cefotaxime and 20 μg/mL hygromycin. The spore content of a single shaking flask was distributed to 3-4 agar plates after collection and washing. Transgenic thalli were observed after 1-2 weeks under white fluorescent light at 23°C.

### Protoplast preparation

Protoplasts were isolated from *M. polymorpha* thalli as previously described_39_, with modifications: thalli were vacuum-infiltrated by 1/2 B5 containing 2%_(w/v)_ Driselase and 6%_(w/v)_ Mannitol for 10 min in a glass beaker, and subsequently incubated in the dark at room temperature for 5 h. The beaker was then gently swirled for 30 sec to aid protoplast release, and the protoplast-containing suspension passed through a Falcon 40 μm cell strainer to remove debris. Protoplasts were isolated from *A. thaliana* as previously described_49_.

### Protoplast encapsulation in microfluidic droplets

Protoplast in the aqueous phase were encapsulated into droplets using a flow-focusing microfluidic device: the protoplast suspension was loaded into a 500 μL Hamilton Gas-tight syringe (Hamilton Robotics, Reno, NV, USA). The fluorinated oil used as the continuous phase (3M Novec 7500 Engineered Fluid with 2.5% PicoSurf 1 surfactant, Sphere Fluidics, Cambridge, UK) was loaded in another syringe and both syringes were connected to the respective inlets of the flow-focusing device (nozzle dimensions: 40 μm x 40 μm x 50 μm) with fine bore polyethylene tubing (ID = 0.38 mm, OD = 1.09 mm, Smiths Medical International, Luton, UK). Using syringe pumps (PHD 2000 Infusion, Harvard Apparatus, Holliston, MA), the two solutions were injected simultaneously in the device. The oil phase was injected at a rate of 500 μL/h and the aqueous phase at a rate of 300 μL/h. The generated droplets were collected, through tubing connected to the outlet, into a syringe.

### Bright-field and fluorescence microscopy

Microdroplet formation was monitored using a Phantom V72 fast camera (Vision Research, Wayne, NJ, USA) mounted on an inverted microscope (IX71, Olympus, Tokyo, Japan). Videos of the encapsulation procedure were captured using the supplied Phantom software. Protoplasts encapsulated in microdroplets were imaged using an inverted microscope (IX71, Olympus, Tokyo, Japan). Chlorophyll fluorescence was excited at 642-682 nm and collected at 603.5-678.5 nm. YFP fluorescence was excited at 488-512 nm and collected at 528.5-555.5 nm.

### On-chip fluorescence measurements and sorting of encapsulated protoplasts

To measure protoplasts fluorescence in each microdroplet, a fixed 491 nm wavelength laser (Cobolt AB, Solna, Sweden) was shaped into a light sheet at 50 mV. The laser was focused through an UPlanFL N 20x microscope objective and directed to the microfluidic chip placed on the stage of an inverted microscope (IX71, Olympus, Tokyo, Japan). Fluorescence detection was carried out by a custom multi-part optical instrument (see Fig. 3A for details). All filters used in this setup were purchased from Semrock (Rochester, NY, USA). Notably, emitted fluorescence was filtered through a 495 nm long-pass filter to eliminate the 491 nm excitation band. Fluorescence was recorded by a PMT (H8249, Hamamatsu Photonics, Shizuoka, Japan), and the data collected was sent to a computer through a DAQ data acquisition card (National Instruments, Austin, TX, USA). The program LabVIEW (National Instruments, Austin, TX, USA) was used to monitor and analyse the data. A microfluidic device was used for sorting YFP-expressing protoplasts in microdroplets (see Fig. 6A): as the microdroplets passed through the objective field of view, they were illuminated by a 491 nm laser. Emitted fluorescence filtered through a 528.5-nm YFP band-pass filter was collected by the PMT and triggered a pulse generator connected to a high-voltage power supply. The resulting electrode pulse deformed YFP-positive microdroplets and targeted them to a small ‘positive’ channel for collection. If the microdroplet was empty or contained protoplast lacking detectable YFP, the PMT sent no signal and the microdroplet passed through the larger ‘negative’ channel.

## Conclusions

We have developed a droplet-based microfluidic platform for high-throughput characterization of plant protoplasts. Our device is capable of quantifying chlorophyll and GFP fluorescence of individual encapsulated cells as a function of genetic circuit activity or in response to environmental stimuli. This workflow allows collection of substantial amounts of biological information from comparatively little plant material. We expect our droplet-based microfluidic platform to be applied for screening of synthetic genetic circuits as well as of mutagenized and enhancer trap lines of a variety of plant species. In the future, we envision a microfluidic workflow composed of on-chip transformation, characterization, and fluorescence-based selection of individual plant cells in preparation of targeted regeneration into whole plants. Combined with libraries of guide RNAs and gene editing tools such as CRISPR-Cas9 nuclease, this workflow promises to greatly accelerate academic and industrial research in modern plant biotechnology.

## Conflicts of interest

There are no conflicts to declare.

## Acknowledgements

We thank the University of Cambridge SynBioFund and the OpenPlant Fund for financial support.

---

**Supplementary Figure S1.**
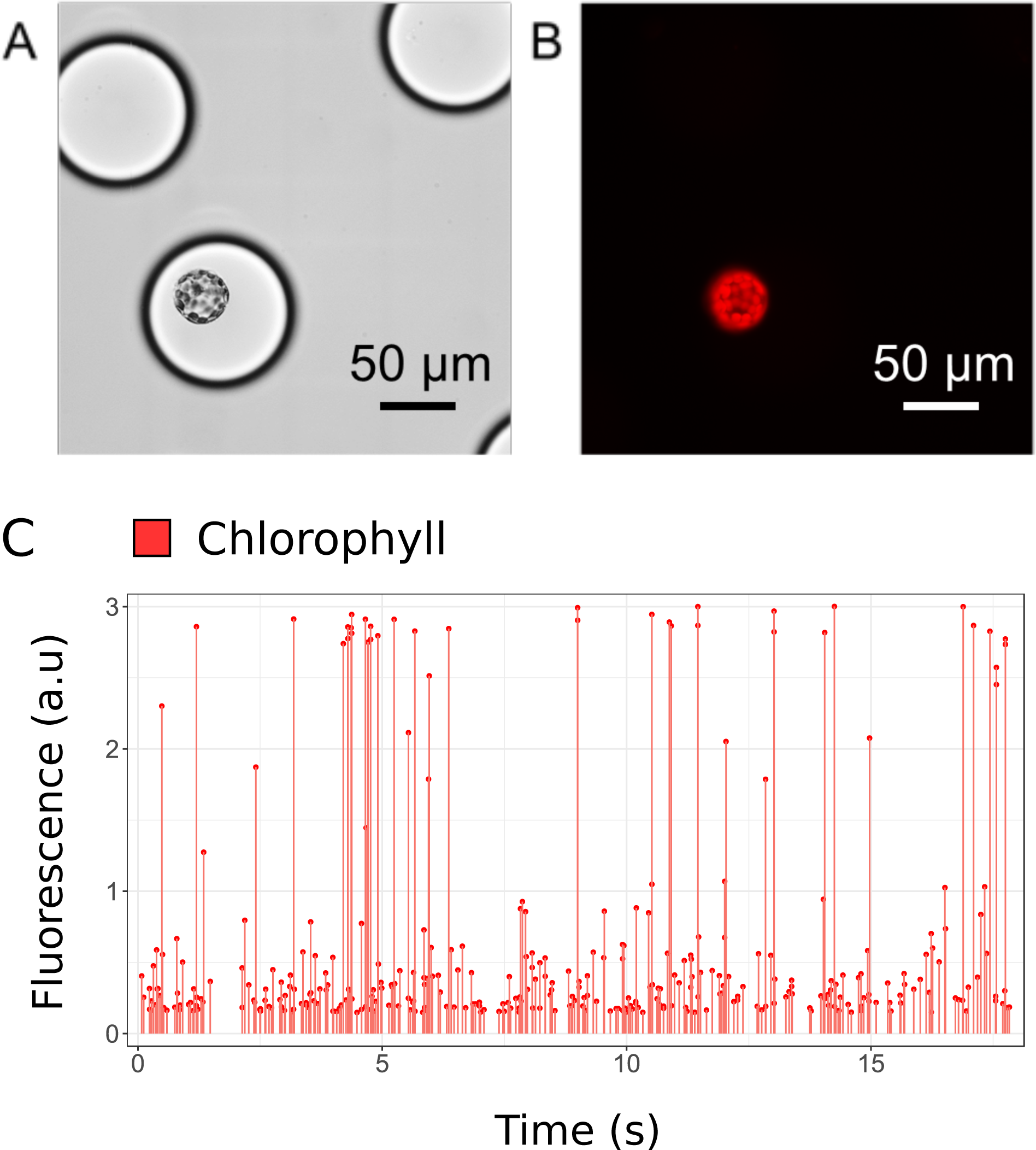
A) Bright field and (B) chlorophyll fluorescence micrographs of individual A. thaliana leaf protoplasts encapsulated in microdroplets. (C) Representative photomultiplier tube (PMT) readout of chlorophyll fluorescence intensity represented as arbitrary fluorescent units (AFU) recorded over 17.5 s. Each line represents an individual encapsulated protoplast.

Supplementary Video S2. Encapsulation of *Arabidopsis thaliana* protoplasts.

Supplementary Video S3. Sorting *of Marchantia polymorpha* protoplasts expressing YFP.

